# Predicting drug combination response surfaces

**DOI:** 10.1101/2024.04.03.586729

**Authors:** Riikka Huusari, Tianduanyi Wang, Sandor Szedmak, Tero Aittokallio, Juho Rousu

## Abstract

Prediction of drug combination responses is a research question of growing importance for cancer and other complex diseases. Current machine learning approaches generally consider predicting either drug combination synergy summaries or single combination dose-response values, which fail to appropriately model the continuous nature of the underlying dose-response combination surface and can lead to inconsistencies when a synergy score or a dose-response matrix is reconstructed from separate predictions. We propose a structured prediction method, comboKR, that directly predicts the drug combination response surface for a drug combination. The method is based on a powerful input-output kernel regression technique and functional modeling of the response surface. As an important part of our approach, we develop a novel normalisation between response surfaces that standardizes the heterogeneous experimental designs used to measure the dose-responses, and thus allows training the method with data measured in different laboratories. Our experiments on two predictive scenarios highlight the suitability of the proposed approach especially in the traditionally challenging setting of predicting combination responses for new drugs not available in the training data.

---

Drug combinations are increasingly used for treatment of various diseases, especially blood cancers and solid tumors [1–3]. In contrast to monotherapies, combination therapies offer the advantages in overcoming intrinsic and acquired resistance in cancer treatment, enhancing drug responses via synthetic lethality, and reducing unwanted side-effects by lowering the dose of individual drugs in the combination [4, 5].

In pre-clinical stages, drug combinations are typically measured in cell lines using dose-response assays. High-throughput screening enables one to measure the responses of pairwise drug combinations at a few selected concentrations of the two drugs (e.g., 5x5 or 8x8 dose-response matrices) [6]. There are several efforts to conduct large-scale drug combination screens in various cancer types [7–9], which have resulted either in fully or partially measured dose-response matrices.

The synergistic effects of drug combinations are further evaluated by summary synergy scores, calculated by the divergence between the measured drug combination responses and the expected non-interaction responses of the single drugs over the full matrix [6]. Multiple synergy models have been proposed to score such divergence based on different assumptions of the expected non-interaction response, for example, the highest single agent (HSA) model [10], Bliss independence model [11] and Loewe additivity model [12]. However, there are disagreements in terms of synergy when using different synergy models, due to large differences in drug concentrations and maximum response values across studies [13].

Since drug combination synergy is evaluated based on multi-dose combination responses, often tested in multiple cancer cell lines with distinct oncogene addictions, large-scale screening of combination effects is required for systematic discovery of new effective and selective combinations. However, to screen pairwise combinations among 100 drugs at 5 different concentrations in 10 cell lines would already require more than one million experimental tests. To speed up such resource and time consuming combination discovery process, machine learning models are needed to narrow down the massive combinatorial search space [14, 15].

A large proportion of current research focuses on the prediction of drug combination synergy rather than dose response values [15–17]. Some research works focus on the prediction of drug combination responses at selected concentrations [18–21]. A major advantage of predicting directly the dose-combination responses is that different synergy models can be applied to the predicted dose-response matrices in post-analysis, making the prediction task independent of a specific synergy metric. One such prediction method is the comboLTR [21], which predicts directly the scalar-valued dose-combination responses by applying latent tensor-based polynomial regression (LTR) [22]. The drug combination dose-response values can be seen as discrete measurements sampled at different concentration values from a continuous dose-response surface where the response value is a function of drugs’ concentrations. Both synergy score prediction and dose-response prediction can be seen as predictions based on the underlying surfaces: if the full continuous dose-response surface is known, the dose-response matrices can be sampled from the known surface at given concentrations, and the synergy scores can then be derived based on the sampled dose-response matrices. Thus the direct prediction of a dose-response surface is more general task than either predicting the single dose-response values, or the synergy scores.

Recently, Rønneberg et al [23] proposed an approach called PIICM, that considers predicting full response surfaces instead of individual dose-combination responses. They proposed a probabilistic prediction model based on Gaussian process regression where the covariance matrices for pairwise drug interactions are parameterised and learned. Notably, their approach can be interpreted as a matrix completion task on the collected response matrix, as the learning system is based on response data only, without using any additional features (cell line or drug features). This means, however, that the approach can not be expected to adapt to settings where a response surface in a test set would contain a drug not seen in training data. This more challenging new drug scenario is important in practical applications, since one cannot assume that that responses of all the drugs of interest would been already tested before either individually or in combination.

In this work, we propose a novel approach for predicting directly the full continuous drug combination dose-response surfaces with a kernel-based structured output prediction model, called comboKR. In contrast to the PIICM model [23], comboKR is based on an inductive learning approach, which predicts the drug combination response surface from input drug features that are easily available from drug databases. Moreover, to overcome the practical issues arising from the heterogeneous experimental design often used in drug combination response measurements, we propose a novel normalisation scheme for comparing drug interaction surfaces. The main goal of the normalisation scheme is to align the dose-response surfaces to be centered around the area where the response changes rapidly as concentrations change. We demonstrate that with comboKR the massive chemical space can be exploited efficiently toward finding novel effective drug combinations beyond the given drug set with known measured responses. To summarize, our contributions are as follows:

- We propose an accurate approach to drug combination response prediction that predicts the full continuous drug combination response surfaces instead of individual dose-response or synergy score values.
- Our surface-valued regression approach takes advantage of a novel normalisation scheme between drug response surfaces that solves issues arising from the heterogeneous experimental designs.
- Important for novel drug combination discovery, our proposed method can be applied to the new drug settings without the need to re-train the model or measure each drug response beforehand.
- In comparison to the baseline comboLTR method [21], and another surface-valued prediction approach, PIICM [23], we show that comboKR achieves superior results especially in the more challenging predictive scenario, where testing is performed on drugs not available in the training stage.

## Overview of ComboKR

### Surface-valued regression

Our focus is on learning to predict the full, continuous drug interaction response surfaces *y* ∈ *𝒴* for the drug pairs (*d*_1_, *d*_2_) ∈ *𝒳* in a given cell line – a challenging structured output prediction problem. This is especially difficult prediction task, since practically each output – a surface – in any such data set is sampled in part or fully from different sets of concentrations than the others, and therefore the dose-combination response matrices are not directly comparable. To solve the problem, we consider adapting an approach that has sometimes been referred to as generalised kernel dependency estimation (KDE) [24] or input output kernel regression (IOKR) [25, 26].

In a nutshell, our proposed model for surface-valued prediction relies on a vector- or function-valued kernel ridge regression (KRR) problem, obtained by mapping the surfaces to the reproducing kernel Hilbert space (RKHS) *ℋ*_*𝒴*_ associated with kernel *k*_*y*_ : *𝒴 × 𝒴 →* ℝ (see Figure 1, (c)). Now, instead of solving directly the structured prediction problem *f* : *𝒳 → 𝒴*, the learning problem has been cast as vector-valued one to learn *g* : *𝒳 → ℋ*_*𝒴*_, after which the prediction to *ℋ*_*𝒴*_ is mapped back to *𝒴* (pre-image problem). Using the closed-form solution for the KRR and assuming a normalised output kernel *k*_*y*_, the final optimisation problem to obtain predictions can be written as

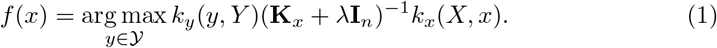

**Fig. 1:**
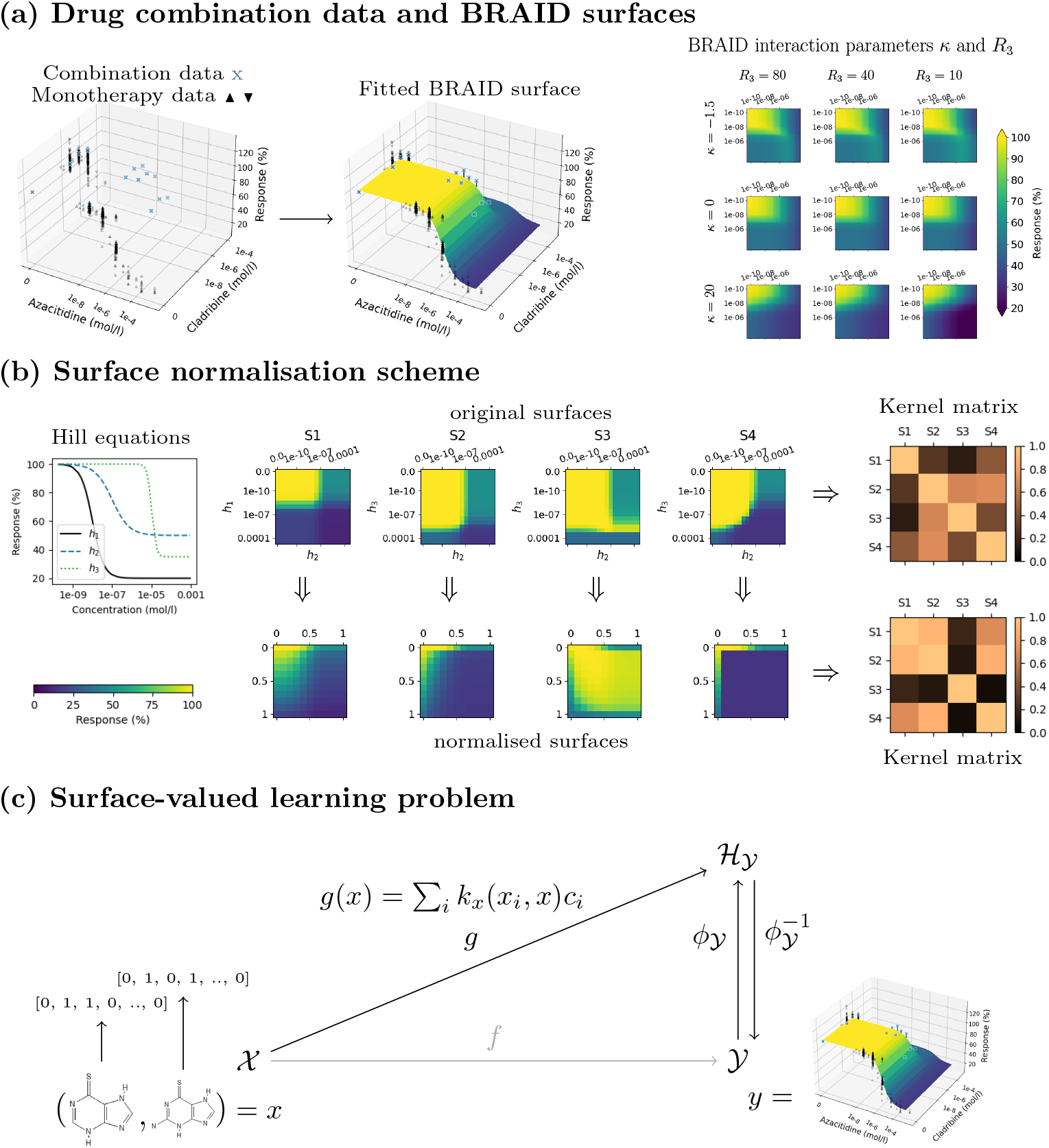
Overview to the proposed approach. (a) The drug combination data and additional monotherapy responses are used to fit a parametric surface on each dose-response matrix. The BRAID surface model uses the Hill equations of the two drugs, as well as two interaction parameters to model different types of response surfaces. (b) Illustration of surface normalisation resulting to different similarities. The surfaces *S*1 and *S*2 are computed with *κ* = 0 (neutral), and *S*2 and *S*4 with *κ* = *−*1.5 (extreme antagonism) and *κ* = 25 (extreme synergism), respectively. (c) Finally, a surface-valued prediction problem is formulated and solved with the output kernel learning -style approach, where the output data is mapped with help of a suitable kernel to RKHS *ℋ*_*𝒴*_.

Here **I**_*n*_ is *n× n* identity matrix, and **K**_*x*_ stands for the *n× n* kernel matrix collecting all values of kernel evaluations between pairs of drugs *k*_*x*_(*x*_*i*_, *x*_*j*_), *i, j* = 1, …, *n* and each *x*_*i*_ stands for a (ordered) pair of drugs, *x*_*i*_ = (*d*_1_, *d*_2_)_*i*_. *λ* is the regularisation parameter of the kernel ridge regression model. The shorthand *k*_*y*_(*y, Y*) refers to the vector [*k*_*y*_(*y, y*_1_), …, *k*_*y*_(*y, y*_*n*_)] with 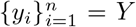 the outputs of the training set; *k*_*x*_(*X, x*) is defined analogously.

Problem (1) is ill-posed in general. When this is the case it is most often solved by restricting the search for maximum value over a candidate set *C*_*y*_ instead of the full output space *𝒴*. Every element in the set is tried out, and the one giving maximum value for (1) is chosen as the final prediction.

### BRAID surfaces

Our learning approach considers full dose-response surfaces as the outputs of the learning problem. Yet, the dose-response data is collected via factorial experimental design, by measuring response values from varying dose-concentrations, resulting in data sets where these surfaces are represented with matrices collecting the measurements. Starting from the famous Loewe additivity [12] and Bliss independence [11], various approaches to modelling drug interactions have been proposed [27–32]. In this work, in order to obtain the continuous form of the surface, we use the BRAID (Bivariate Response to Additive Interacting Doses) drug interaction model [31] that builds on the Hill equation [33, 34] and is motivated by the Loewe additivity principle. We fit this function to each of the drug pair combinations. The model is intuitive, as it uses the Hill equation parameters of the two drugs, in addition to two interaction parameters. More concretely, the function depends on the following parameters:

- Four response parameters: the baseline response in absence of drugs, *R*_0_; the maximal responses of drugs 1 and 2 as *R*_1_ and *R*_2_; and optionally also the maximal combination response *R*_3_^1^
- Hill equation slope parameters for both drugs: *τ*_1_ and *τ*_2_
- Half maximal effective concentration (EC_50_) for the two drugs: *EC*_1_ and *EC*_2_
- The interaction parameter *κ* ∈] *−* 2, *∞*]: *κ <* 0, = 0, *>* 0 for antagonism, additivity or synergy, respectively (as illustrated in Figure 1 (a)).

Now, the BRAID function for drugs 1 and 2 applied in dose concentrations *c*_1_ and *c*_2_ is written as:

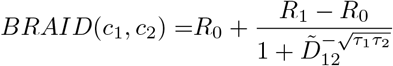

where

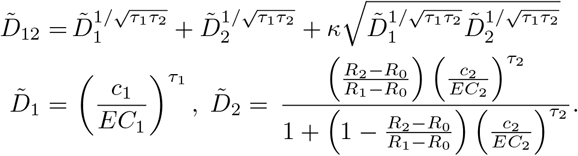

We note that in this formula it is assumed that one of the drugs is the weaker one, and the other stronger. In the practical implementation^2^ either of the drugs could be the stronger one, resulting to a slightly more complicated formula. The implementation provides methods to fit the BRAID functions to data.

Due to the scarcity of the combination data (for example only 3*×*3 measurements for each combination in the NCI-ALMANAC dataset), to aid with the fitting process we additionally include the more abundant monotherapy data available from other sources. We illustrate this inFigure 1 (a).

### Kernels between surfaces

Our surface-valued prediction approach relies on having a kernel, *k*_*y*_, defined between the drug interaction surfaces. We choose to use the Gaussian (or RBF) kernel, defined as 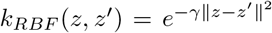 for some vectors *z* and *z*^*′*^. At the first glance, it might seem attractive to use the (vectorised) dose-response matrices directly available in drug combination datasets in this kernel. However, even if those matrices were of same size, most often the drug doses used to measure drug combination responses are widely different between any two surfaces. Thus, these measurements cannot be directly compared to each other. We instead assume that we have functions *S*_*i*_ parameterising the drug interaction surfaces available – in our work we obtain these from the BRAID interaction model. We can now fix a s et o f c oncentrations f or a ll drugs, *C* = [*c*_1_, *c*_2_, …, *c*_*N*_ ], and compare any two surfaces in a RBF kernel with

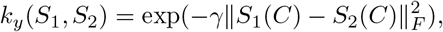

where *S*_1_ and *S*_2_ are response surface functions that are queried from both drug 1 and drug 2 with concentrations *C*, resulting in two *N × N* matrices in the Frobenius norm evaluation in the kernel.

However, this straightforward approach still has limitations. Different drugs often have different effective concentrations, meaning that two combination surfaces expressing very similar interaction profile (e.g. s imilar level o f s ynergy), might b e s hifted in relation to each other such, so that the kernel evaluation results in non-intuitive values. To overcome this issue, we propose normalisation over the dose-concentration values to effective concentration range [ 0, 1 ] using the Hill equation, where 0 corresponds to concentrations that have no effect t o t he c ell growth, 0 .5 indicates E C_50_ concentration, and 1 corresponds to concentrations with maximal response. We denote kernel acting on normalised surfaces as 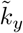.

With this normalisation, problems related to shifted surfaces are decreased, and the kernel 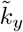 shows more realistically the differences in interaction pattern (i.e. i f the combination is synergistic or antagonistic). We illustrate this difference between the two choices of kernels in Figure 1 (b), where surfaces of three types of interactions (*S*2, *S*3 and *S*4) are compared to baseline surface *S*1. The three surfaces differ from *S*1 by one of the Hill functions (*h*_3_ instead of *h*_1_) and the interaction parameter *κ*. Intuitively, the surfaces *S*1 and *S*2 are closest together in a sense, that their drug combination interaction effect is very similar (*κ* = 0). H owever, t he H ill equations and especially their EC_50_ values for the first d rug are different (equation *h*_1_ vs *h*_3_), which has resulted in the two surfaces being “shifted” w.r.t each other. The Frobenius norms between the original surfaces are in all three cases large, and *S*1 is judged most similar to *S*3. The normalised comparison can take the shifting into account, and *S*2 is therefore judged to be most similar to *S*1.

## Results

In this section, we computationally validate our proposed surface-valued kernel regression model for drug combination response prediction, comboKR. We compare it to a scalar-valued dose-response prediction baseline, comboLTR [21], and to another surface-valued prediction model derived from the PIICM presented in [23] – we call this PIICM^*∗*^ (see Methods section for details). Due to the computational cost of comboKR arising from the use of kernel matrices, we consider all the cell lines as independent data sets, and train and test on them separately – a challenging setting that often arises in real-world personalized medicine applications. For comboKR we consider both original and normalised output kernels, and denote as “comboKR raw” or “comboKR r.” the version with the original, simpler kernel.

The results presented here are obtained from two representative predictive scenarios: new drug and new combo. New combo refers to the case, when the test set consists of new drug combinations – however all the drugs are available in the training set as parts of other combinations. New drug is more realistic, but also more challenging scenario, where the drug combinations in the test set always contain a drug that has not been present in any of the combinations in the training set.

### Overview of the results

We present the overall Pearson correlations averaged over the cell lines between the predicted values and ground truth measurements, as well as between the Bliss and Loewe synergies calculated from them (Figure 2). For the response values, we also display density plots in Figure 3. In the easier new combo scenario, the scalar-valued prediction approach, ComboLTR, slighly outperforms the surface-valued ones. However, in the more challenging new drug scenario, the situation reverses, and the surface-valued approaches mostly outperform ComboLTR. In this more challenging setting, comboKR has a clear advantage over the other models, especially when using the concentration normalised surface kernels. PIICM^*∗*^ predictions in the new drug scenario has problems for some predictions, for which the algorithm does not converge from the values used in initialisation (see supplementary material). The vertical anomalies in the density plots of the surface-valued models originate from the failures in fitting the BRAID model accurately to the data (see Figure 9 in the Methods section).

**Fig. 2:**
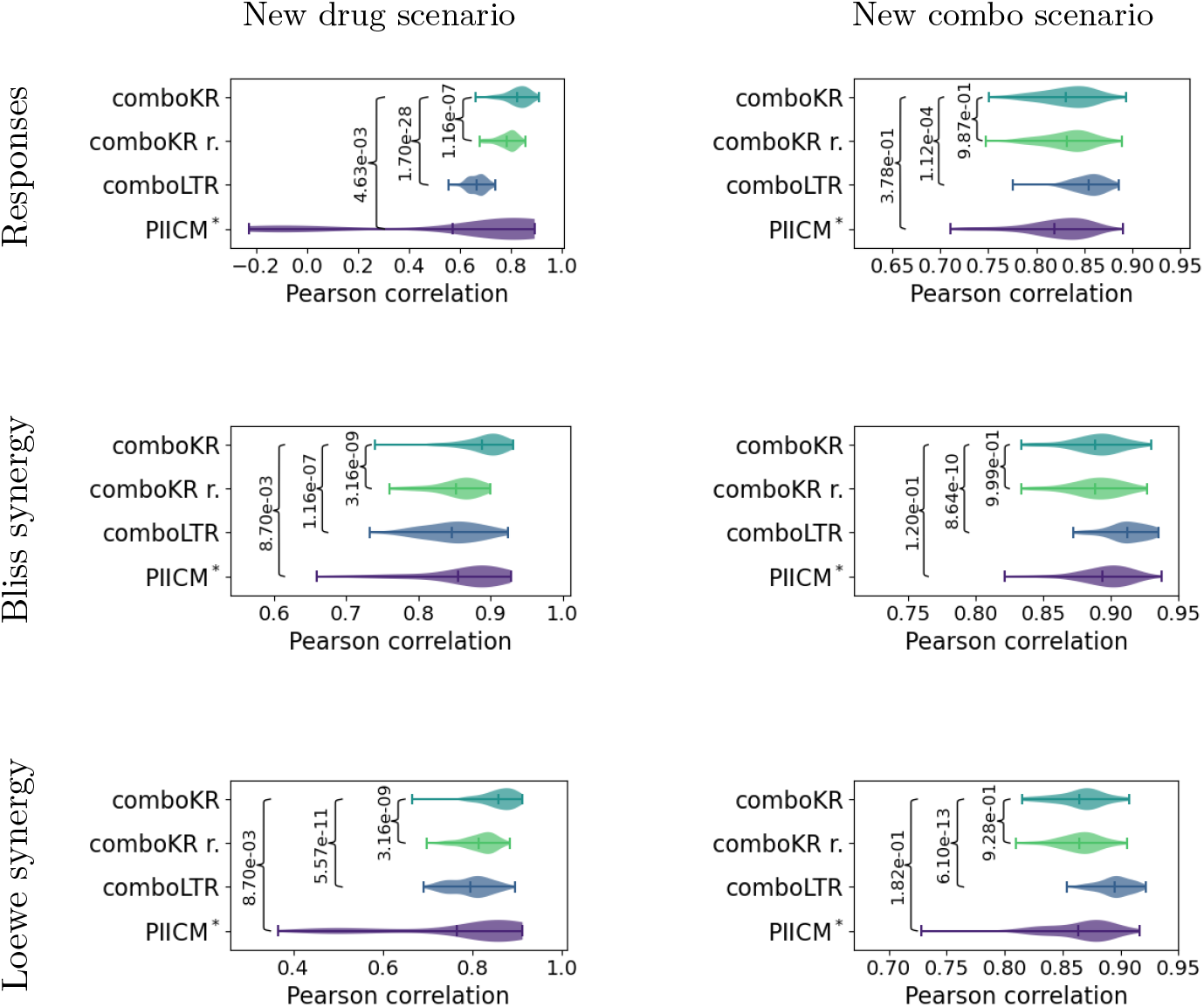
Violin plots of Pearson correlations over the 60 cell lines in the two prediction settings. The vertical lines in the plots highlight the mean and the extrema. First row shows correlation between original and predicted combination responses. Additionally, synergy scores (Bliss and Loewe) were calculated from both ground truth measurements as well as for the predictions. The two other rows show correlations of these synergy scores. Statistical p-values from two-sided Kolmogorov-Smirnov test are shown for all the competing methods compared to ComboKR (full pairwise results are available in Supplementary material).

**Fig. 3:**
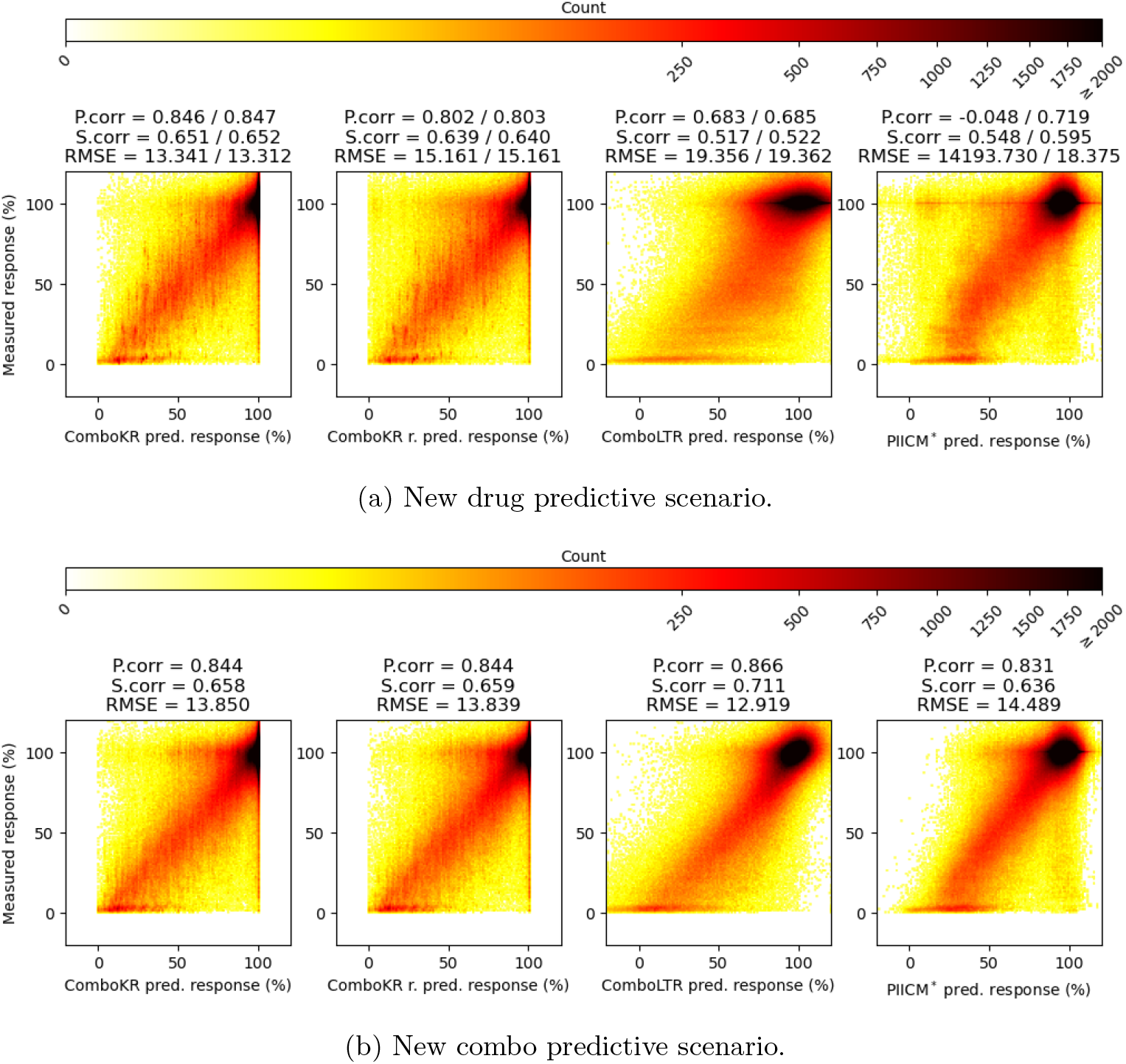
Density plots of predictions and ground truth measurements over all cell lines for the two predictive scenarios. The titles of the plots indicate the Pearson correlation, Spearman correlation and root mean squared error (RMSE) between the predicted and the measured response values. In the new drug predictive scenario, the second values reported are over the subset of responses where PIICM^*∗*^ predictions were not outliers (see supplementary material).

It can be seen from Figure 3, that the surface-valued models can offer improvements over the scalar-valued model in predicting the extreme response values that are more rare in the data. The comboLTR model focuses on the predictions of the more abundant, higher response values, and easily overshoots the predictions of the lower response values. While this is easier to see in the new drug predictive scenario, the same behaviour is already present in the new combo setting. The other surface-valued method, PIICM^*∗*^, predicts the extreme response values better than comboLTR, but not as well as the comboKR approaches.

In supplementary material, we show additional results for the correlations with the fitted BRAID surfaces, as well as full pairwise results on the statistical significance of the differences between results displayed in Figure 2.

### Performance over tissue and drug combination types

The NCI-ALMANAC’s cell lines originate from nine tissue types. Similarly, the tested drugs belong to three drug groups (chemotherapy, targeted and other). Details of these groupings can be found from the supplementary material.

We investigated and compared in more detail the performance of the comboKR and comboLTR methods on the different tissue types and different drug type combinations in the two predictive settings (Figure 4). As before, comboLTR method performs slightly better than the comboKR approaches in the easier predictive scenario. However, in the new drug scenario, differences between comboKR and comboLTR is much larger, favouring the comboKR approaches. The comboKR variants perform relatively similarly in the easier setting, but especially when looking at the different cell lines, the normalised kernel comboKR outperforms the usual ones.

**Fig. 4:**
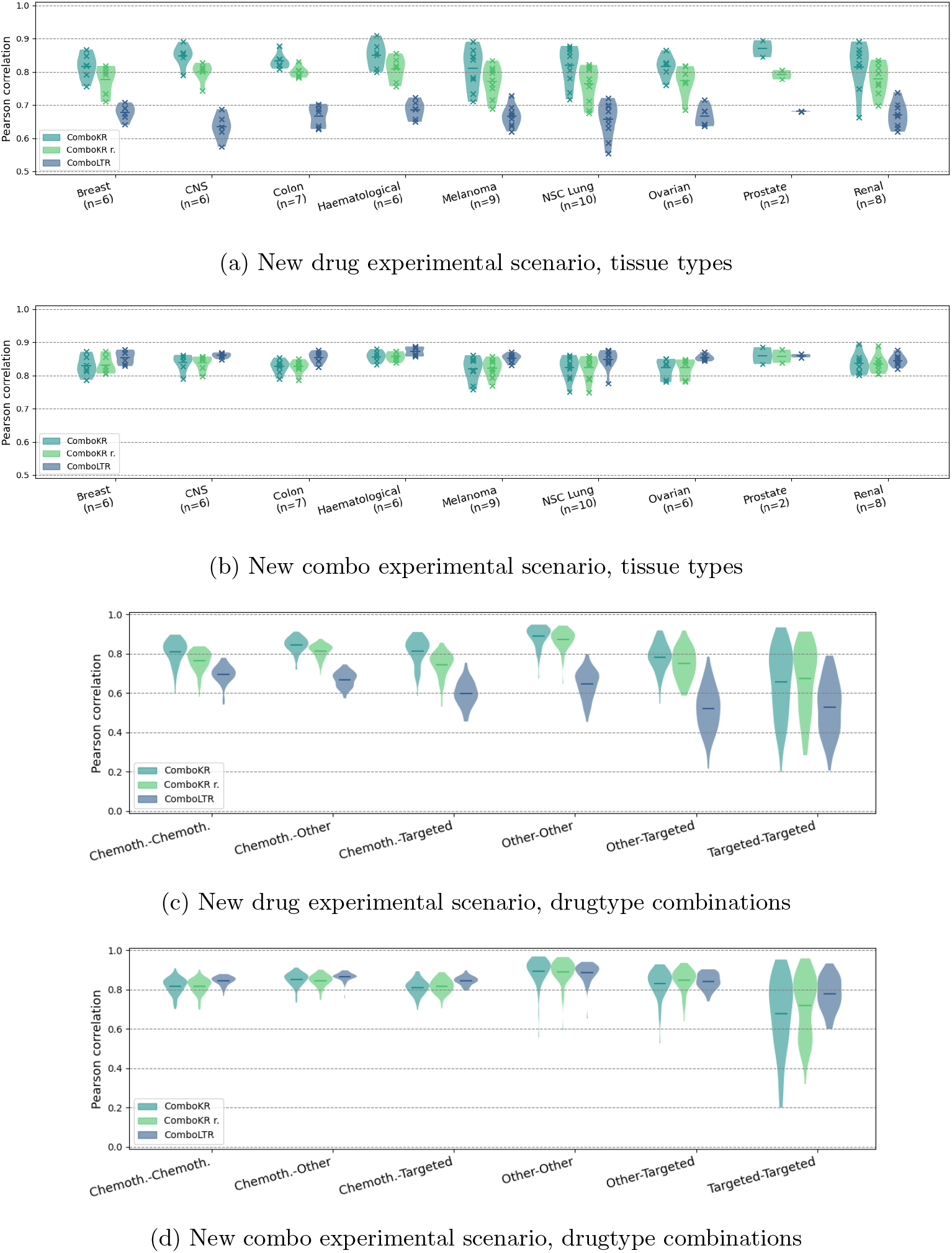
Distributions of Pearson correlations of the drug-dose response prediction on the different tissue and drug type combinations (see supplementary material for details) in the two predictive scenarios, for the comboKR variants and comboLTR. The vertical lines in the plots highlight the mean. The violins with fewer than ten elements (i.e. results over tissue types) indicate the number of elements in parenthesis of the x axis label, and additionally include also the individual results, marked with crosses – drugtype combination results are over all the 60 cell lines.

### Performance of predicting individual surfaces

The results so far have focused on overall performance of the models. Here, we show in Table 1 the results of pairwise comparing individual surface predictions. The table reports the average percentages (averaged over the cell lines) how often a method achieved a better prediction for the surfaces in the test set, w.r.t mean squared error, compared to the other methods (the comparison is always pairwise).

**Table 1:** Pairwise comparison of the methods. The table reports average percentages over all 60 cell lines how often did the method on row give better prediction for a surface, w.r.t mean squared error, than the method on column; rows with high values (darker colouring; similarly columns with low values with lighter colouring) indicate better performance for the method. The two comboKR versions with original and normalised surface kernels use the same candidate sets, and so their predictions might be identical, resulting in ties in the rankings. Tie counts are not included in the tables, so cells marked with asterisk do not add to 100%.

| % | c.KR r. | c.KR | c.LTR | PIICM* |
| --- | --- | --- | --- | --- |
| c.KR r. | | 34.9±4.6* | 68.3±6.4 | 53.7±7.3 |
| c.KR | 57.2±5.4* | | 76.3±6.1 | 64.9±6.0 |
| c.LTR | 31.7±6.4 | 23.7±6.1 | | 31.0±13.2 |
| PIICM* | 46.3±7.3 | 35.1±6.0 | 69.0±13.2 | |

(b) New combo scenario. \*comboKR versions resulted in ties at rate $14.7\% \pm 2.6\%$ .
| % | c.KR r. | c.KR | c.LTR | PIICM* |
| --- | --- | --- | --- | --- |
| c.KR r. | | 41.9±2.9 * | 45.6±4.4 | 49.3±4.4 |
| c.KR | 43.4±3.5 * | | 48.3±5.2 | 51.6±4.8 |
| c.LTR | 54.4±4.4 | 51.7±5.2 | | 52.5±6.4 |
| PIICM* | 50.7±4.4 | 48.4±4.8 | 47.5±6.4 | |

Similarly to the previous results, also here it can be observed that the differences between the methods are small in the easier new combo predictive scenario (Table 1b). The new drug predictive scenario (Table 1a) highlights again the benefits of the surface-valued methods: comboLTR obtains most often the worst prediction for a surface, even though PIICM^*∗*^ did neither result in a good prediction for some of the surfaces. The results again highlight how using the normalised surface kernels out-performs the basic comboKR approach – also the percentage of identical predictions between the two versions is almost half the amount of the new combo setting.

We further highlight the advantage of our comboKR against comboLTR in Figure 5. ComboLTR predicts accurately only those concentrations measured in the drug response assay, separately, which can lead to predictions that do not follow a smooth interaction pattern. In contrast, comboKR predicts a full continuous interaction surface, from which one can sample any dose-concentration pairs to obtain dose-specific predictions or summary synergy scores.

**Fig. 5:**
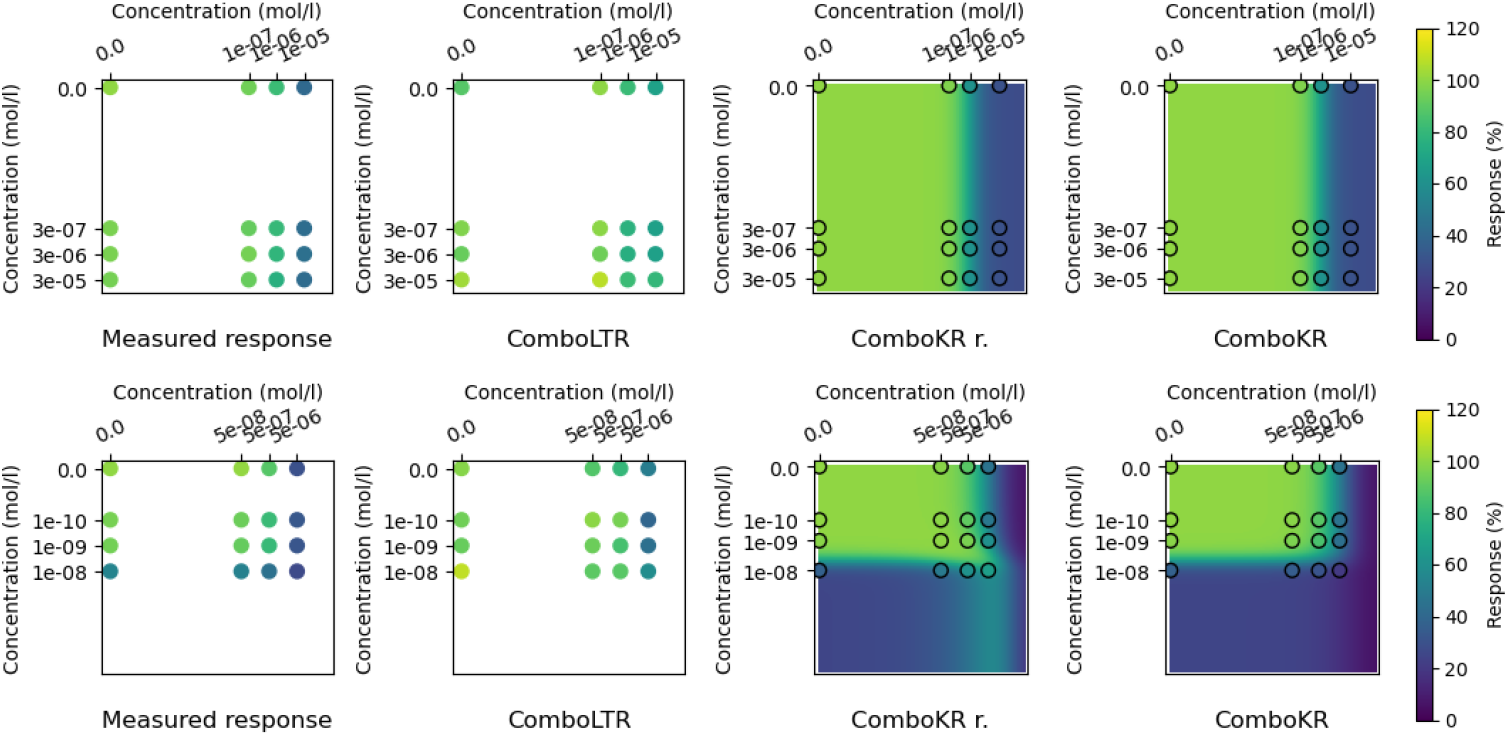
Measured responses and example predictions from comboLTR and the two comboKR versions in the new drug setting. Top row shows combination of drugs thioguanine (id 752, horizontal axis) and lenalidomide (id 747972, on vertical axis), and bottom row of drugs triethylenemelamine (id 9706, on horizontal axis) and SN-38 (id 673596, on vertical axis), both of cell line 786-0.

## Discussion

In this work, we have investigated the drug combination response prediction problem from the point of view of predicting entire drug combination surfaces, instead of predicting individual response values. We propose an approach based on kernel methods, which when combined with a novel surface normalisation scheme overcomes issues arising from the heterogeneous experimental design used to measure the data. We show that casting the drug combination response prediction as a structured prediction learning problem can improve predictive performance especially in the traditionally challenging experimental settings. Namely, our method shows great promise especially in the new drug scenario, providing the opportunity to find promising drug combinations that go beyond the limited set of drugs in the training set.

To explore the suitability of our proposed surface-valued learning approach, we performed computational experiments on the NCI-ALMANAC dataset. Of the two predictive scenarios investigated in our experiments, the proposed surface-based approach achieves better predictive performance especially in the more challenging new drugs scenario. Even the more straightforward surface-valued prediction approach outperformed the baseline comboLTR method, but especially the novel concentration normalisation provies significant improvements in this challemging, yet practical setting. In personalized medicine studies, either focusing on indivudal cell lines or patient samples, one cannot assume that each drug has already been tested in combination with other drugs in often limited training datasets. In addition to outperforming other approaches, the suitability of our proposed comboKR in the new drugs scenario is also computationally practical: unlike with the traditional methods, the proposed model does not need to be re-trained to obtain predictions when a new set of drugs is introduced to the test set. Moreover, as illustrated in Figure 5, the comboKR predicts a full continuous drug interaction surface, instead of individual values that might not conform to a smooth interaction pattern. Again, this gives a practical advantage: it is easier based on predicted surface to experimentally validate the synergy between two drugs by using the surface to determine relevant drug concentrations. Surface sampling can be also used to suggest doses for experimental testing with highest likelihood for revealing synergy between two drugs.

Additionally, we observed that the surface-valued methods in general were better suited for predicting extreme response values (Figure 3). This was most clearly observed in the new drugs setting. Even in this case, our proposed comboKR approaches outperformed the other surface-valued method, obtaining more accurate predictions on the extreme response values. The extreme responses are often most informative for identifying synergistic (or antagonistic) interaction between two drugs, so their prediction is critically important for drug combination discovery.

Our experiments give promising results for using structured prediction approach in the drug combination response prediction, motivating future research. It will be important to investigate ways to make the ComboKR model more scalable on big high-throughput screening data consisting of millions of data points. The obvious bottleneck for applications to multiple cell lines is the sampling of the large kernel matrix on inputs, consisting of two Kronecker products: *K*_*c*_ *⊗ K*_*d*_ *⊗ K*_*d*_. In addition to efficient algorithms and paralellisation strategies, for example, kernel approximations could be investigated to speed up the computations. Another avenue to pursue would be to follow [36], and investigate the scalability using or generalising the proposed Kronecker product vec-trick.

## Methods

### The NCI-ALMANAC drug combination data

In this study, we consider the drug combination dataset NCI-ALMANAC [7]. It provides systematic screening of drug combinations among 104 FDA-approved anticancer drugs on the 60 NCI-60 human tumor cell lines covering 9 different tissue types (see Supplementary material). In this dataset, drugs have been screened at either 5 or 3 concentrations, resulting in 5*×*3 or 3*×*3 drug combination dose-response matrices. To have consistent dose response matrix size, only 3*×*3 dose response matrices were kept for further modeling (311527 combinations), as the number of 5*×*3 dose-response matrices is much fewer than 3*×*3 matrices. Monotherapy responses at combination dose concentrations were also included in the NCI-ALMANAC dataset for the 104 drugs, where single drug responses were measured at various concentrations. Different concentration values were determined for different drugs [7]. Thus, each drug combination response surface is represented by the measured responses by a 4*×*4 matrix containing both combo- and mono-responses. When duplicate measurements of the same drug combination on the same cell line appeared, the median of measurements was taken.

The NCI-ALMANAC data has been collected with standard factorial experimental design. The 3-by-3 (or 4*×*4) dose response matrices described above are independent from each other in a sense, that the dose levels between experiments (matrices) used to collect the data are not the same. Notably, even if two response matrices both consider the same drug as one factor, the concentrations used in measurements might differ between them. Figure 6 shows that here is a large amount of different concentration combinations in the NCI-ALMANAC data. On average, any given dose combination is found from only 0.44% of the drug response matrices. The most common concentration combination is present at 12.4% of the matrices (the full distribution is shown in Supplementary Material). It is very rare that for any two surfaces, all nine concentration combinations would match. Thus, directly comparing any two dose-response matrices in the dataset can very rarely be done.

**Fig. 6:**
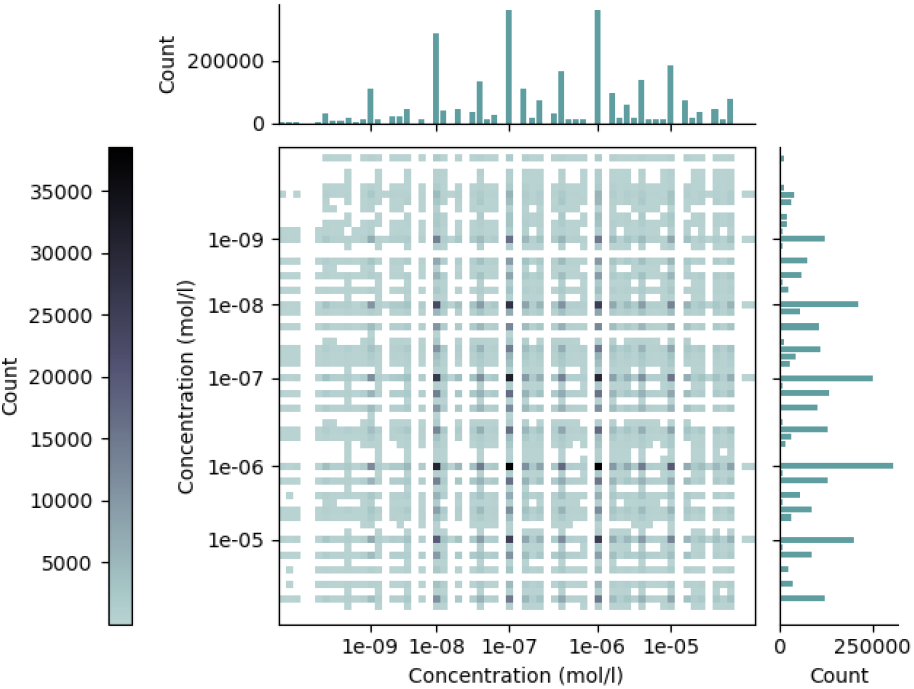
The counts of drug doses and their combinations in the NCI-ALMANAC dataset.

NCI-ALMANAC reported two endpoints calculated differently by using time zero measurement as reference or not [37]. For the percent growth of cells with time zero as reference (”PercentGrowth” as reported in the dataset), the responses range from -100 to 100. However, for this endpoint, the calculation processes are different when the percent growth of test cells are lower or higher than the time zero measurement of cell growth. Whereas for the percent growth of cells using only control values (”PercentGrowthNoTZ” as reported in the dataset) as reference, the response calculation is consistent and ranges from 0 to 100. Thus, for simplicity and consistency of data, the endpoint denoted as “PercentGrowthNoTZ” was used as the response value. If there were missing values for the endpoint, it was calculated from the raw measurements reported in the dataset.

Our proposed comboKR, as well as the compared PIICM method, rely on accurate surface modeling in training stage. Since the drug combination response data only contains three monotherapy measurements for both drugs, the NCI-60 single drug monotherapy response data where typically a drug is measured at multiple concentrations were also integrated as part of the surface model fit procedure, in order to help improve the model performance with better estimate of monotherapy dose response functions.

To train the machine learning models to predict the dose-response values, we consider the commonly used 166-bit 2D structure MACCS molecular fingerprints as the input data.

### The ComboKR model

Our proposed ComboKR model is an adaptation of an approach that has sometimes been referred to as generalised kernel dependency estimation (KDE) [24] or input output kernel regression (IOKR) [25, 26].

In this approach, operator-valued kernels (OvKs) are used to solve a regression problem to a reproducing kernel Hilbert space (RKHS) *ℋ*_*𝒴*_ induced by a traditional scalar-valued kernel *k*_*y*_ defined for the output data in *𝒴*. OvKs are associated with vector-valued reproducing kernel Hilbert spaces (vv-RKHSs), containing functions that map the data to the vector-valued (or function-valued) output space. Thus, they are a natural choice to solve for the function *g* : *𝒳 → ℋ*_*𝒴*_.

In this context, the operator-valued kernel is a function *𝒦* : *𝒳 × 𝒳 → L*(*ℋ*_*𝒴*_) in which *L*(*ℋ*_*𝒴*_) denotes the set of linear operators from *ℋ*_*𝒴*_ to *ℋ*_*𝒴*_ - if the output space *ℋ*_*𝒴*_ is finite-dimensional, i.e. *ℋ*_*𝒴*_ = ℝ^*p*^, then *ℒ* (*ℋ*_*𝒴*_) = ℝ^*p×p*^. In practise, the most common operator-valued kernel to use is the separable (or decomposable) kernel, which can be written as *𝒦* (*x, z*) = *k*_*x*_(*x, z*)**T** with *k*_*x*_ : *𝒳 × 𝒳 →* ℝ being a traditional scalar-valued kernel on the input data, and **T** ∈ *ℒ* (*ℋ*_*𝒴*_), which in turn is often chosen to be the identity.

Operator-valued kernels generalise the usual scalar-valued kernels, notably also the representer theorem. Thus, the solution to the regularised learning problem considered in the IOKR,

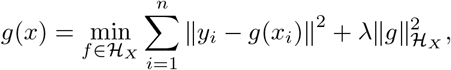

where *ℋ*_*X*_ denotes the vv-RKHS associated with *𝒦*, can be written as

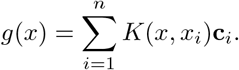

Here **c**_*i*_ ∈ *ℋ*_*𝒴*_ are the multipliers to be learned. Like in the usual scalar-valued case, the closed-form solution

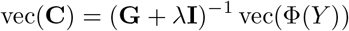

can be obtained, in which Φ(*Y*) = [*ϕ*_*y*_(*y*_1_), *ϕ*_*y*_(*y*_2_), …, *ϕ*_*y*_(*y*_*n*_)] and **C** = [**c**_1_, **c**_2_, …, **c**_*n*_], both of size *d × n* if the size of feature space associated with kernel *k*_*y*_ is denoted with *d*. **G** is the *nd × nd* operator-valued kernel matrix with block-wise structure. Now,

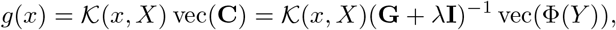

where *𝒦* (*x, X*) = [*𝒦* (*x, x*_1_), …, *𝒦* (*x, x*_*n*_)]

After solving for *g*, the final p redictions c an b e o btained f rom t he pre-image problem

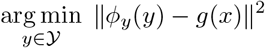

or from

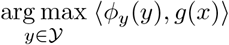

if one assumes that the output kernel *k*_*y*_ is normalised, i.e. *k*_*y*_(*y, y*) = 1 *∀y* ∈ *𝒴*. It is now possible to take advantage of the form of the separable kernel matrix, and the Kronecker product vec-trick, to obtain the final optimisation problem

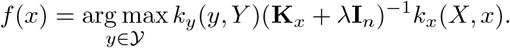

Here **I**_*n*_ is *n × n* identity matrix, and **K**_*x*_ stands for the *n × n* kernel matrix 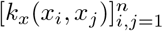. The shorthand *k*_*y*_(*y, Y*) with 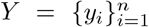 the training set outputs, refers to the vector [*k*_*y*_(*y, y*_1_), …, *k*_*y*_(*y, y*_*n*_)]; *k*_*x*_(*X, x*) is defined analogously. The pseudocode of the approach can be found in Algorithm 1.

#### Algorithm 1

The comboKR approach

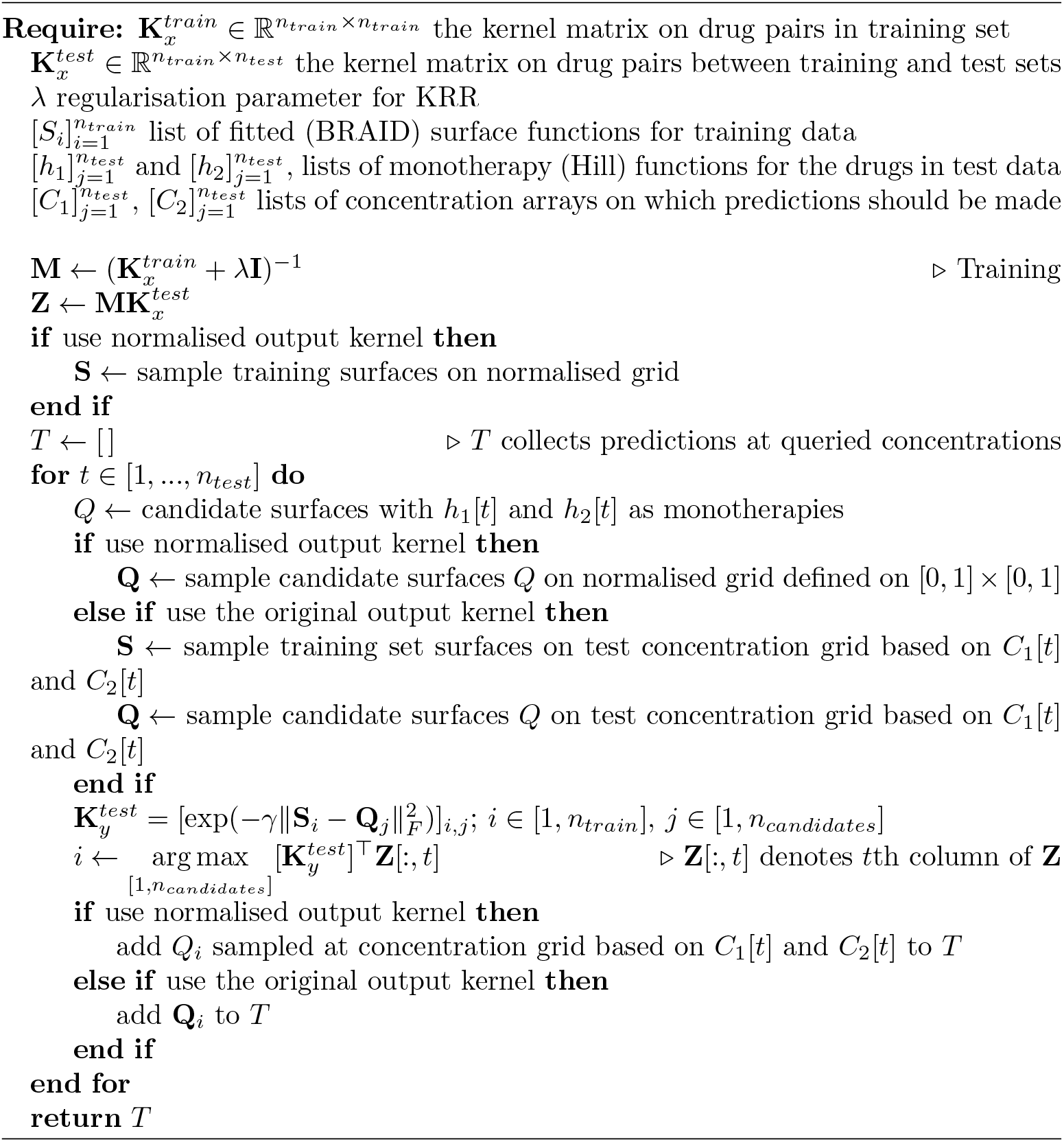

In this work we choose *k*_*x*_ to be defined as 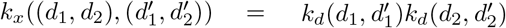 with *k*_*d*_ acting on the MACCS fingerprints to be Tanimoto kernel [38]. As discussed, for the output surfaces we consider the RBF kernel, that conforms to the assumption of having the output kernel being normalised. We use the parameter 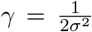 with *s* equal to mean of distances between the surfaces in the training set.

### Concentration normalisation to obtain a normalised surface kernel

The main idea of our proposed surface normalisation scheme is to standardise the concentration measurements across the different drugs, so that all the concentrations are in same range, and the surface comparisons can be made more easily. To this end, we map the dose-concentrations with help of Hill equations, individually for each drug in each cell line, to the range [0, 1], where 0 intuitively stands for “no effect”, and 1 stands for “maximal effect” (see Figure 7). More concretely, the concentration normalisation transformation *CN* can be written as

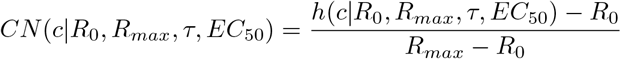

in which *c* is the original concentration, and *h* is the Hill equation with parameters *R*_0_, *R*_*max*_, *τ* and *EC*_50_ (i.e. baseline response, maximal response, slope parameter and half maximal effective concentration). The normalisation formula uses the same *R*_0_ and *R*_*max*_ as the Hill equation.

**Fig. 7:**
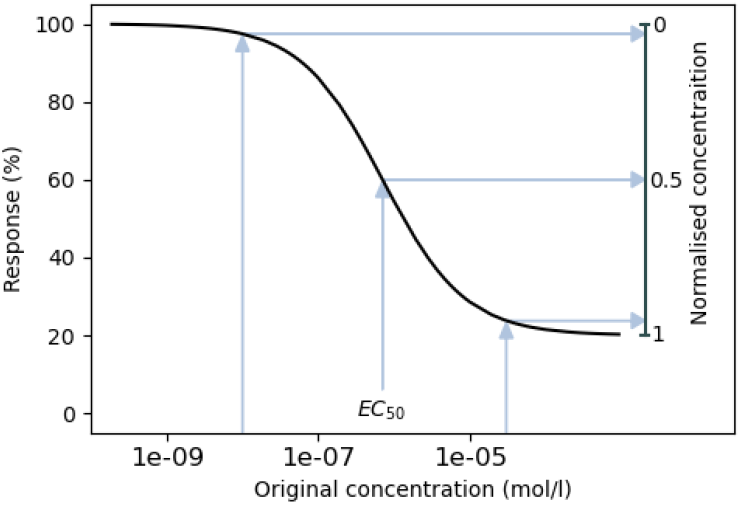
Illustration of the concentration normalisation procedure to [0, 1] range. The concentrations too low to elicit a response are mapped to zero, EC_50_ concentration is mapped to 0.5, and concentrations close to maximal effect are mapped close to 1.

After the concentration normalisation, the values [0, 1] can be seen to correspond to each other over all the different drugs – unlike in non-normalised case where a concentration might yield a very different response from two different drugs. With all drug concentrations normalised, comparing different surfaces is easy in a common grid of values from [0, 1]*×*[0, 1]. Due to the nature of the normalisation scheme, it can be expected that this common drug grid in all surfaces will focus especially on areas where the potential synergy or antagonism can best be captured (as illustrated in second part of Figure 1).

### Experimental setup

We compare our comboKR^3^ to two baselines: comboLTR^4^ [21, 22] for predicting individual response values, and PIICM^5^ [23] as another surface-valued prediction method based on Gaussian processes. With our comboKR, we consider both the original and normalised surfaces in output kernel computations: the former case is denoted by comboKR r.

ComboLTR is a polynomial regression model for scalar-valued prediction that exploits higher order interactions between the views in predictions. As input for a prediction, it takes the two drugs and their concentrations. As with our comboKR, the drugs are represented by the MACCS fingerprints. The comboLTR method represents the concentrations as one-hot encoded vectors.

PIICM is a surface-valued approach based on Gaussian process regression. With PIICM we use the same BRAID surfaces as the baseline models as in our comboKR. Like in [23], we consider a subset of concentrations to represent the surfaces since the method is computationally too heavy to run with full set of concentrations in the data (over 60 unique dose values), and the BRAID surfaces are sampled at these concentrations as training data to the model. As the concentrations are assumed to be normalised to [0, 1] interval, we consider the same normalisation scheme for PIIMC, as we use to obtain our normalised kernel evaluations.

Yet, we observed that the memory requirements for the PIICM method to run in our setting were infeasible, as the number of drugs considered is relatively large. In order to run the method, we forced it to consider a simplified form of the drug combination covariance matrix – we call this PIICM^*∗*^. While this might put the method at disadvantage, in practice we observe that in the new combo setting its performance is close to comboLTR, which is close to results obtained in [23]. Moreover, we observed in small-scale experiments that our modification performs comparably to, or even slightly improves, the original parametrisation (see details in Supplementary Material).

The PIICM^*∗*^ outputs predictions on the chosen concentration grid. To obtain the final PIICM^*∗*^ predictions, we interpolate with Nadaraya–Watson kernel regression from this set of concentrations to the concentrations the test surfaces are measured at.

The parameters for the models are chosen with cross-validation, taking into account the specifics of the predictive scenario in each split. For our comboKR, *λ* is chosen from `1e-2, 5e-2, 1e-1, 5e-1`, while in comboLTR, the order is considered 3 or 5, and rank 10 or 20. PIICM^*∗*^ rank is cross-validated over 5, 25, 50, 75 and 100. In all models the training data is “doubled”, i.e. both orders of drug pairings are included separately in the training set.

### Predictive scenarios

We consider predictions in two scenarios:

- **New drug:** one of the drugs in the combination queried at the test stage has not been seen in any combination during the training stage. Monotherapy responses of the novel drug are assumed to be available during the training stage.
- **New combo:** the drug combination queried at the test stage has not been seen in training; however, both single drugs may have been seen in the combinations encountered during training.

In the new combo setting, the data in each cell line is divided evenly into five folds, of which one is used as test fold, and the remaining folds are used in training and validation. In the new drug setting, the test fold contains all surfaces where either drug is one of ten randomly chosen as the new drug. The four validation folds are obtained similarly among the rest of the data used in the training, always with five randomly chosen drugs. Clearly, the new combo setting is the easier of the two prediction scenarios. We note that in both scenarios, no drug-dose response values of a surface queried at test point are available in training. The task is to predict all the combination responses, instead of filling missing values inside a matrix with known entries.

In the more challenging new drugs scenario, the drug features and monotherapy responses for the novel drug are assumed to be available already in the training stage. For most of the standard models, including comboLTR baseline, it means that at any time a new drug is to be included in testing, the model would need to be retrained with the additional information. This can be very costly. An advantage of our proposed comboKR method is that this kind of re-training is not needed, and the new monotherapy response data can only be introduced to the model during the test stage, when the candidate surfaces are being built. In this scenario, the relevant monotherapy responses of the new drugs in test set are included to augment the comboLTR training set. Similarly, in PIICM, those monotherapy responses are included in the response matrix the method takes as input.

We note that the PIICM method [23] is based on Gaussian processes, modelling the drug interaction surfaces and without considering any drug features when making predictions. Thus, it is not expected to be able to generalise to data outside of the training set in the new drug scenario, further than predicting some generic interactions derived from the other drugs.

### Obtaining the drug interaction surfaces

As a preprocessing step for surface-based approaches (PIICM and our proposed comboKR), we fit the BRAID drug interaction surfaces to the available combo data with the synergy package [35]. Due to the scarcity of the monotherapy responses in the NCI-ALMANAC combination data, the available monotherapy responses were used to augment the combo data. Hill equations fitted to the monotherapy data are used in building the normalised output kernels. Overall, the average Pearson correlation of the fitted BRAID surfaces sampled at the measurement positions to the ground truth combination measurements was 0.9037 *±* 0.0169 averaged over the cell lines, 0.908 over all of them (see Figure 8 for density plot).

**Fig. 8:**
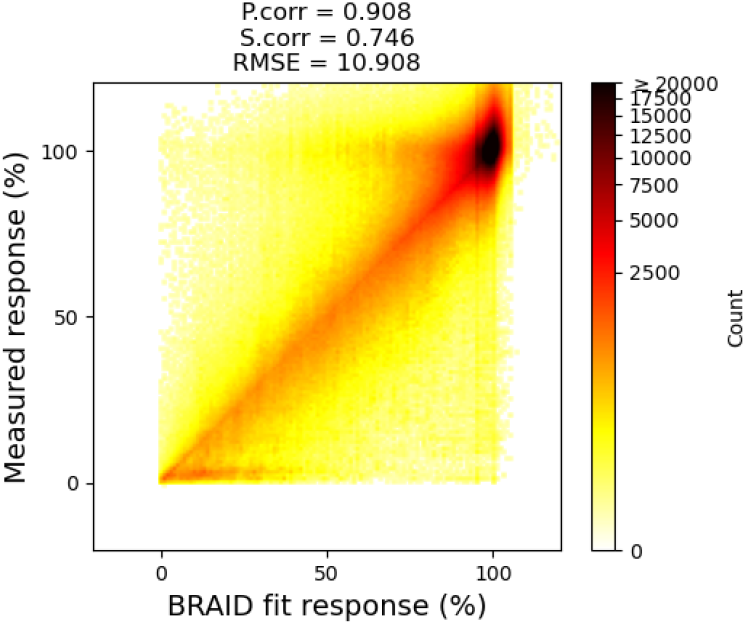
Density plot of the measured responses compared to the responses sampled from the fitted BRAID surface models at the same concentrations, over the full NCI-ALMANAC data.

**Fig. 9:**
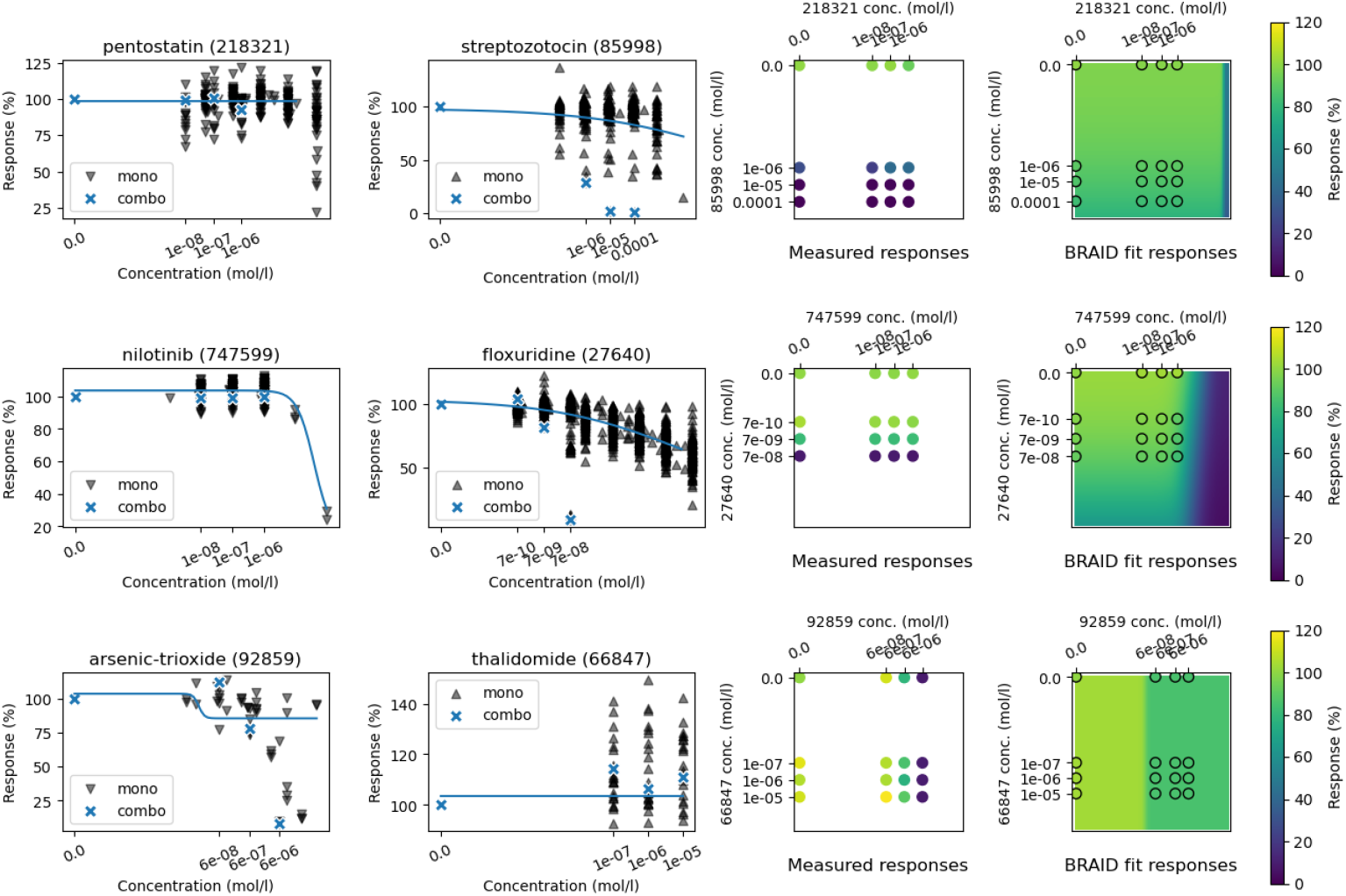
Examples of BRAID surfaces not fit well to the NCI-ALMANAC dataset. On top: cell line HL-60(TB), drugs pentostatin (id 218321) and streptozotocin (id 85998). Middle: cell line NCI-H226, drugs nilotinib (id 747599) and floxuridine (id 27640). Bottom: cell line OVCAR-3, drugs arsenic-trioxide (id 92859) and thalidomide (id 66847). The two first plots in each row show the NCI-60 monotherapy responses in black, with blue crosses highlighting the NCI-ALMANAC monotherapy response values, while the curve displays the responses from the Hill equation from BRAID model. Third column displays the full combo data matrix, and final column shows the fitted BRAID surface.

It can be expected that large amount of of the discrepancies are due to surface model filtering out noise from the measurements. However, some errors in fitting are due to the conflicts between NCI-ALMANAC and NCI-60 datasets. Examples of this are provided in Figure 9.

### Candidate surfaces for comboKR

The simplest way to solve the pre-image problem (i.e. finding the best *y* for prediction *g*(*x*) in *ℋ*_*𝒴*_) for the comboKR problem 4, is to consider a candidate set where all elements are tested out, and the one giving the highest score is selected as the prediction. In both of the predictive scenarios, we assume that the monotherapies of the drugs are available. Thus, we can easily create relevant surface candidates with the surface model by only varying the parameters *κ* and *E*_*max*_ that are related to the behaviour of the drug combination. In experiments, the candidates are generated by sampling *κ* from [-1.99, -1.5, -1, -0.5, -0.1, 0.01, 0.1, 0.5, 1, 2, 10, 25, 50] (to capture various surface interaction profiles), and *E*_*max*_ from around the maximum values of the individual drug responses.

### Performance metrics

The test sets in both predictive settings consist of drug-dose interaction surfaces that have been sampled at various differing 4 *×* 4 grids. The predictions at those concentrations were obtained as follows: in comboLTR they are directly predicted, in PIICM they are interpolated from the predictions in the normalised grid, and in our comboKR they are sampled from the predicted BRAID surface. Two synergy scores (Bliss and Loewe) are calculated based on these ground truth and predicted 4 *×* 4 matrices.

We report Pearson correlations of the predictions (both for raw responses and summary synergy scores) calculated for each cell line separately. Within a cell line, both the predictions and ground truth labels are vectorised, and the correlation is calculated between the two vectors. The same procedure is used when investigating different tissue types and drug combination types: all the ground truth and predicted responses within the group in a cell line are vectorised to compute the Pearson correlation.

Furthermore, we compare all the methods to each other on the level of individual surfaces, by reporting how often in the test set each of the methods obtained the best prediction for a given surface with respect to the mean squared error – Pearson correlation not being applicable if the matrix of values that is sampled from the predicted surface is constant. This happens most often in modified PIICM, but also sometimes in both comboKR versions.

## Data availability

The NCI-60 and NCI-ALMANAC datasets [7] used are publicly available at https://wiki.nci.nih.gov/display/NCIDTPdata/NCI-60+Growth+Inhibition+Data and https://wiki.nci.nih.gov/display/NCIDTPdata/NCI-ALMANAC.

## Code availability

The code for the comboKR method is available at https://github.com/aalto-ics-kepaco/comboKR. For the competing methods, we used codes made available for comboLTR at https://github.com/aalto-ics-kepaco/ComboLTR and adapted PIICM from https://github.com/ltronneb/PIICM/. The software for the BRAID surfaces is available from https://github.com/djwooten/synergy/.

## Footnotes

1 The *R*_3_ parameter is not present in original BRAID model introduced in [31], but it is present in the implementation [35].

2 https://github.com/djwooten/synergy

3 Available at https://github.com/aalto-ics-kepaco/comboKR

4 https://github.com/aalto-ics-kepaco/ComboLTR

5 https://github.com/ltronneb/PIICM/

