## Supplementary material for "Predicting drug combination response surfaces"

### Predicting drug combination response surfaces Supplementary Material

#### NCI-ALMANAC dataset

Figure 1 shows the amounts of the $3 \times 3$ dose-response matrices for each cell line in the NCI-ALMANAC dataset. Figure 2 displays the amounts of the monotherapy responses from NCI-60 dataset that were used.

Figure 3 shows the full distribution of in how many surfaces concentration combinations are present. Majority of the dose combinations can be found in very few surfaces.

The tissue types in the NCI-ALMANAC dataset and lists of cell lines belonging to them are listed in Table 1. Similarly, the drugs in the dataset and their types are listed in Table 2.

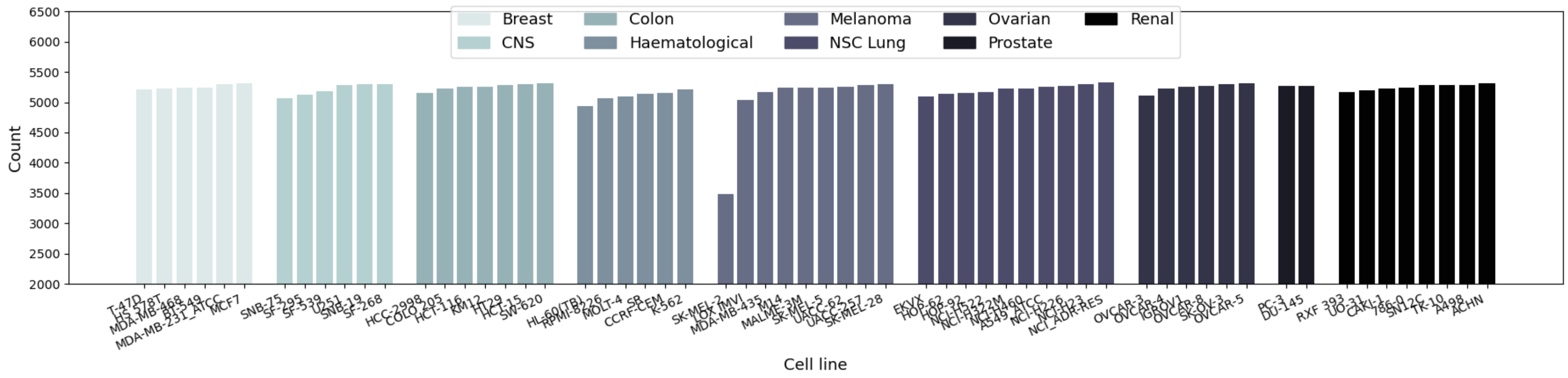


Figure 1: Counts of the $3 \times 3$ dose-response measurement matrices (i.e. dose-response surfaces) in the processed NCI-ALMANAC dataset per cell line.

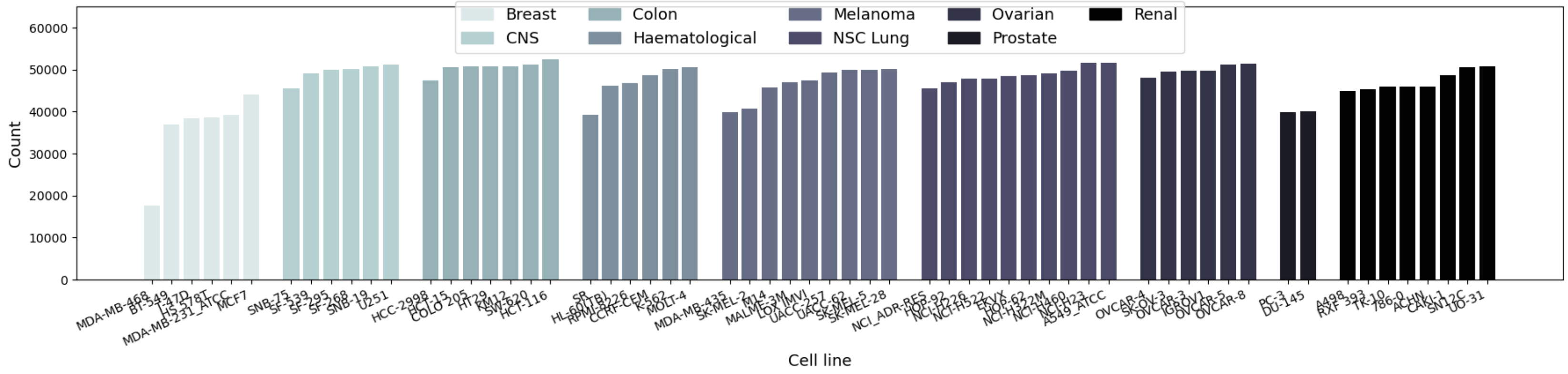


Figure 2: Counts of individual dose-response measurements from the NCI-60 dataset used to assist in fitting the BRAID surfaces.

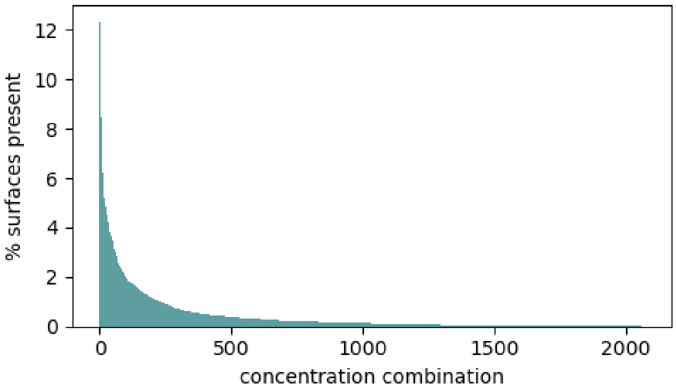


Figure 3: Distribution on how many surfaces a concentration combination is present in (as percentage), in the NCI-ALMANAC drug combo dataset. The dose combinations are sorted in descending order w.r.t the amount of surfaces they are found in.

Table 1: The tissue types and the cell lines belonging to them.

| Tissue type | Count | Cell line names |
|---|---|---|
| Breast | 6 | BT-549, HS 578T, MCF7, MDA-MB-231/ATCC, MDA-MB-468, T-47D |
| CNS | 6 | SF-268, SF-295, SF-539, SNB-19, SNB-75, U251 |
| Colon | 7 | COLO 205, HCC-2998, HCT-116, HCT-15, HT29, KM12, SW-620 |
| Haematological | 6 | CCRF-CEM, HL-60(TB), K-562, MOLT-4, RPMI-8226, SR |
| Melanoma | 9 | LOX IMVI, M14, MALME-3M, MDA-MB-435, SK-MEL-2, SK-MEL-28, SK-MEL-5, UACC-257, UACC-62 |
| NSC Lung | 10 | A549/ATCC, EKVX, HOP-62, HOP-92, NCI-H226, NCI-H23, NCI-H322M, NCI-H460, NCI-H522, NCI/ADR-RES |
| Ovarian | 6 | IGROV1, OVCAR-3, OVCAR-4, OVCAR-5, OVCAR-8, SK-OV-3 |
| Prostate | 2 | DU-145, PC-3 |
| Renal | 8 | 786-0, A498, ACHN, CAKI-1, RXF 393, SN12C, TK-10, UO-31 |

Table 2: The drug types

| Drug id | Drug name | Drug type | Drug id | Drug name | Drug type |
|---|---|---|---|---|---|
| 740 | methotrexate | Chemotherapy | 246131 | valrubicin | Chemotherapy |
| 750 | busulfan | Chemotherapy | 256439 | idarubicin | Not Used |
| 752 | thioguanine | Chemotherapy | 256942 | epirubicin | Not Used |
| 755 | mercaptopurine | Chemotherapy | 266046 | oxaliplatin | Chemotherapy |
| 762 | Mechlorethamine hydrochloride | Chemotherapy | 279836 | mitoxantrone | Chemotherapy |
| 1390 | allopurinol | Other | 296961 | amifostine | Chemotherapy |
| 3053 | dactinomycin | Chemotherapy | 362856 | temozolomide | Chemotherapy |
| 3088 | chlorambucil | Chemotherapy | 369100 | imiquimod | Other |
| 6396 | thiotepa | Chemotherapy | 409962 | carmustine | Chemotherapy |
| 8806 | melphalan | Chemotherapy | 606869 | clofarabine | Chemotherapy |
| 9706 | Triethylenemelamine | Chemotherapy | 608210 | vinorelbine | Chemotherapy |
| 13875 | altretamine | Chemotherapy | 609699 | topotecan | Chemotherapy |
| 14229 | mepacrine | Chemotherapy | 613327 | gemcitabine | Chemotherapy |
| 18509 | 5-aminolevulinic-acid | Other | 628503 | docetaxel | Chemotherapy |
| 19893 | 5-fluorouracil | Chemotherapy | 673596 | SN-38 | Chemotherapy |
| 24559 | plicamycin | Chemotherapy | 681239 | bortezomib | Chemotherapy |
| 25154 | pipobroman | Chemotherapy | 686673 | nelarabine | Chemotherapy |
| 26271 | cyclophosphamide | Chemotherapy | 698037 | pemetrexed | Chemotherapy |
| 26980 | mitomycin-C | Chemotherapy | 701852 | vorinostat | Chemotherapy |
| 27640 | floxuridine | Chemotherapy | 702294 | estramustine-phosphate | Chemotherapy |
| 32065 | hydroxyurea | Chemotherapy | 707389 | eribulin | Not Used |
| 34462 | uracil-mustard | Chemotherapy | 712807 | capecitabine | Chemotherapy |
| 38721 | mitotane | Chemotherapy | 713563 | exemestane | Other |
| 45388 | dacarbazine | Chemotherapy | 715055 | gefitinib | Targeted therapy |
| 45923 | methoxsalen | Chemotherapy | 718781 | erlotinib | Targeted therapy |
| 49842 | vinblastine | Chemotherapy | 719276 | fulvestrant | Other |
| 63878 | cytarabine | Chemotherapy | 719344 | anastrozole | Chemotherapy |
| 66847 | thalidomide | Other | 719345 | letrozole | Other |
| 67574 | vincristine | Chemotherapy | 719627 | celecoxib | Chemotherapy |
| 71423 | Megestrol acetate | Other | 721517 | zoledronic-acid | Other |
| 77213 | procarbazine | Chemotherapy | 732517 | dasatinib | Targeted therapy |
| 79037 | lomustine | Chemotherapy | 733504 | everolimus | Targeted therapy |
| 82151 | daunorubicin | Chemotherapy | 737754 | pazopanib | Targeted therapy |
| 85998 | streptozotocin | Chemotherapy | 743414 | imatinib | Targeted therapy |
| 92859 | arsenic-trioxide | Chemotherapy | 745750 | lapatinib | Targeted therapy |
| 102816 | azacitidine | Chemotherapy | 747599 | nilotinib | Targeted therapy |
| 105014 | cladribine | Chemotherapy | 747971 | sorafenib | Targeted therapy |
| 109724 | ifosfamide | Chemotherapy | 747972 | lenalidomide | Other |
| 118218 | fludarabine | Chemotherapy | 747973 | ixabepilone | Other |
| 119875 | cisplatin | Chemotherapy | 747974 | raloxifene | Other |
| 122758 | tretinoin | Other | 749226 | abiraterone | Chemotherapy |
| 122819 | teniposide | Chemotherapy | 750690 | sunitinib | Targeted therapy |
| 123127 | doxorubicin | Chemotherapy | 753082 | vemurafenib | Targeted therapy |
| 125066 | bleomycin | Chemotherapy | 754143 | romidepsin | Other |
| 125973 | paclitaxel | Chemotherapy | 754230 | pralatrexate | Chemotherapy |
| 127716 | decitabine | Chemotherapy | 755986 | vismodegib | Targeted therapy |
| 138783 | bendamustine | Chemotherapy | 756645 | crizotinib | Targeted therapy |
| 141540 | etoposide | Chemotherapy | 757441 | axitinib | Targeted therapy |
| 169780 | dexrazoxane | Other | 760766 | vandetanib | Targeted therapy |
| 180973 | tamoxifen | Other | 761431 | vemurafenib | Targeted therapy |
| 218321 | pentostatin | Chemotherapy | 761432 | cabazitaxel | Chemotherapy |
| 226080 | sirolimus | Other | 763371 | ruxolitinib | Targeted therapy |
| 241240 | carboplatin | Chemotherapy | | | |

#### PIICM modification

In order to assess the suitability of our PIICM modification (PIICM*), we performed a small scale experimental comparison to the original parametrisation. To this end, we subsampled the data in cell line "786-0" (chosen as it was first in alphabetical order), and chose either every third or every fourth drug from alphabetical order to be included – choosing every second already resulted to memory errors with original PIICM. The data was divided into training, validation and testing according to the new combo scenario. The partition sizes are shown in Table 3.

Table 3: Data sizes in PIICM modification test runs.

| Setting | #drugs | #surfaces | $n_{tr}$ | $n_{val}$ | $n_{tst}$ |
|---|---|---|---|---|---|
| Every 3rd | 35 | 569 | 341 | 113 | 115 |
| Every 4th | 26 | 296 | 177 | 59 | 60 |

PIICM takes as input a matrix $\mathbf{Y}$ collecting the response data, where a row corresponds to concentrations of the two drugs, and column to drug combination. While one can use as the columns just the drug combinations in training and test sets, in our modification we enumerate all combinations, to make the kronecker product structure of the covariance matrix make sense. Thus, the $\mathbf{Y}$ is larger in our modification. We compare to original PIICM parametrisation with both $\mathbf{Y}$ structures, and denote with PIICM$^\dagger$ the version using $\mathbf{Y}$ as in our modification.

The different ranks used in PIICM cross-validation were selected from [10, 20, 25, 50, 75, 100, 150, 200, 250, 300, 400, 500]:

- In our modification, the rank is at most equal to number of drugs; thus, here the ranks were [10, 20, 25]
- In original parametrisation the drug rank is at most number of surfaces in the data. Thus, the ranks 300, 400 and 500 are only applicable to the setting where every third drug was selected.

For comparison, comboKR was cross-validated with $\lambda$ values in [1e-5, 1e-4, 1e-3, 1e-2, 1e-1].

The results obtained on the test set are displayed in Figure 4. It is clear that our modification does not negatively impact the PIICM results - in fact, it seems to offer a clear improvement compared to the original parametrisation. Running times of the final training-test cycle are displayed in Table 4.

Table 4: Running time of final training-test cycle

| Method | running time (h:min:s) | |
|---|---|---|
| | every 3rd | every 4th |
| comboKR (both) | 0:00:09.236669 | 0:00:02.603092 |
| PIICM* | 0:00:38.839474 | 0:00:24.557288 |
| PIICM$^\dagger$ | *memory error* | 1:07:09.720523 |
| PIICM | 0:27:10.222010 | 0:15:39.983792 |

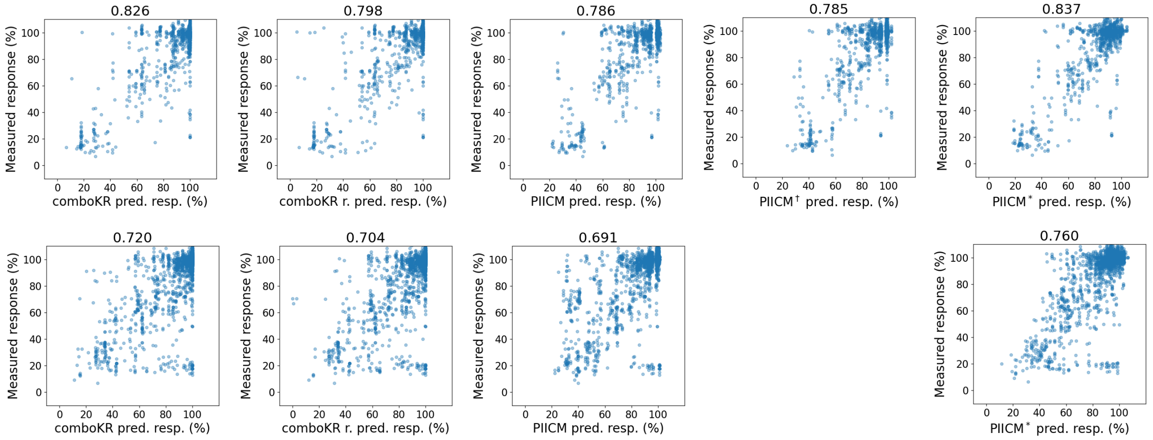


Figure 4: Results of the small-scale experiments of PIICM modification comparison. The methods shown in the columns are comboKR, comboKR with normalised kernel, PIICM with original parametrisation, PIICM$^{\dagger}$ with our modified input, and PIICM$^{*}$ with our modified input and parametrisation. Titles indicate the Pearson correlation to the measured responses. Top row: experimental setting "every fourth", bottom row: "every third". Memory error occurred for PIICM$^{\dagger}$ with original parametrisation in the larger experiment.

#### Supplementary results

##### Scatter plots

We show complete scatter plot of the responses in the new drug predictive scenario in Figure 5, where also the PIICM* predictions not converged from the initialisation are included.

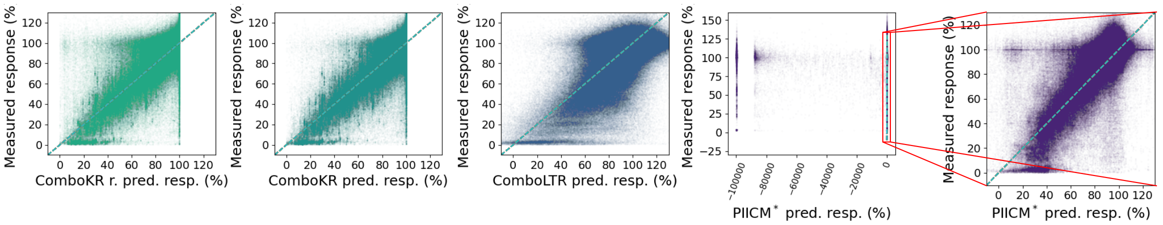


Figure 5: Scatter plots of predicted and measured responses over all cell lines in the new drug predictive scenario. The red rectangle in the left PIICM* prediction scatter plot highlights the area that is shown in the right PIICM* prediction scatter plot.

##### Pearson correlation tables

Table 5 displays mean and standard deviation of Pearson correlations, for both predicted response values, and the Bliss and Loewe synergies calculated from them. As ground truth, both the original measurements and the BRAID surfaces fit to the data and sampled at corresponding concentrations are considered. As ComboKR and PIICM* rely on the fitted surfaces instead of the raw measurements, it is natural that the performance with respect to BRAID is higher.

##### p-values for the results

Fig. 2 in the main manuscript presented selected p-values of the pairwise comparison of the results. Here, we present all the p-values from two-sample Kolmogorov–Smirnov test in Table 6. In the new drug predictive scenario, differences between comboKR and other methods are always statistically significant.

##### Tissue type and drug combination type comparison with PIICM

Figure 6 shows comparison between methods within tissue and drug combination types as shown in the main paper for comboKR variants and comboLTR, here including also PIICM* results.

Table 5: Pearson correlations between the ground truth and predicted responses, as well as between Bliss and Loewe synergy scores of the ground truth and predicted responses. The results are averaged over all cell lines (± standard deviation). The performance at test stage is measured to both original ground truth measurements, as well as to BRAID surfaces sampled at those concentrations.

(a) New combo scenario

| | Responses | |
|---|---|---|
| | Measurements | BRAID |
| comboKR | 0.831 ± 0.028 | 0.923 ± 0.017 |
| comboKR n. | 0.831 ± 0.030 | 0.922 ± 0.022 |
| comboLTR | 0.854 ± 0.018 | 0.855 ± 0.017 |
| PIICM* | 0.818 ± 0.038 | 0.896 ± 0.037 |

| | Bliss synergy scores | |
|---|---|---|
| | Measurements | BRAID |
| comboKR | 0.888 ± 0.021 | 0.936 ± 0.016 |
| comboKR n. | 0.888 ± 0.021 | 0.935 ± 0.018 |
| comboLTR | 0.912 ± 0.015 | 0.914 ± 0.013 |
| PIICM* | 0.894 ± 0.023 | 0.934 ± 0.019 |

| | Loewe synergy scores | |
|---|---|---|
| | Measurements | BRAID |
| comboKR | 0.865 ± 0.020 | 0.920 ± 0.017 |
| comboKR n. | 0.865 ± 0.022 | 0.919 ± 0.021 |
| comboLTR | 0.895 ± 0.015 | 0.894 ± 0.013 |
| PIICM* | 0.864 ± 0.032 | 0.908 ± 0.033 |

(b) New drug scenario.

| | Responses | |
|---|---|---|
| | Groundtruth | BRAID |
| comboKR | 0.782 ± 0.043 | 0.866 ± 0.032 |
| comboKR n. | 0.825 ± 0.049 | 0.912 ± 0.036 |
| comboLTR | 0.665 ± 0.037 | 0.675 ± 0.038 |
| PIICM* | 0.572 ± 0.400 | 0.627 ± 0.434 |

| | Bliss synergy scores | |
|---|---|---|
| | Groundtruth | BRAID |
| comboKR | 0.852 ± 0.034 | 0.895 ± 0.029 |
| comboKR n. | 0.888 ± 0.034 | 0.931 ± 0.029 |
| comboLTR | 0.846 ± 0.042 | 0.857 ± 0.035 |
| PIICM* | 0.856 ± 0.064 | 0.894 ± 0.066 |

| | Loewe synergy scores | |
|---|---|---|
| | Groundtruth | BRAID |
| comboKR | 0.813 ± 0.039 | 0.865 ± 0.036 |
| comboKR n. | 0.857 ± 0.042 | 0.910 ± 0.037 |
| comboLTR | 0.795 ± 0.046 | 0.804 ± 0.039 |
| PIICM* | 0.764 ± 0.158 | 0.805 ± 0.168 |

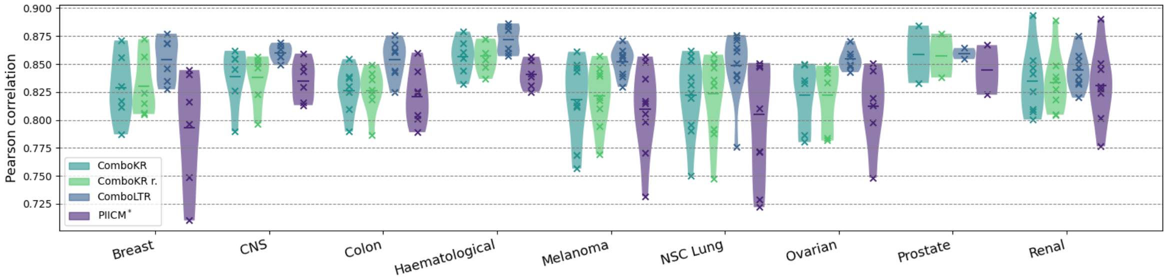


(a) New combo experimental scenario, tissue types

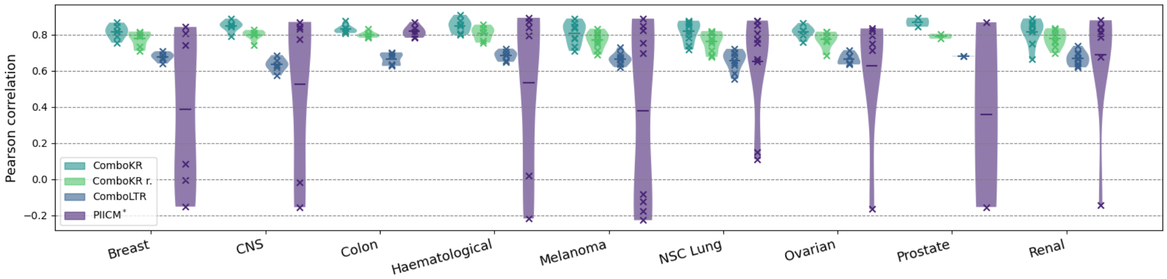


(b) New drug experimental scenario, tissue types

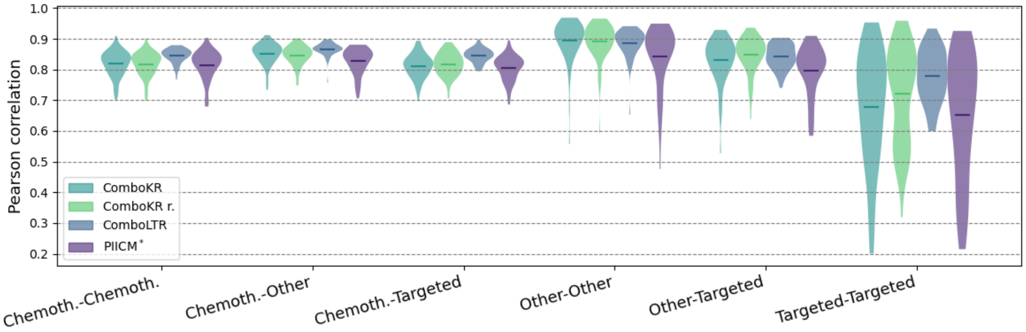


(c) New combo experimental scenario, drugtype combinations

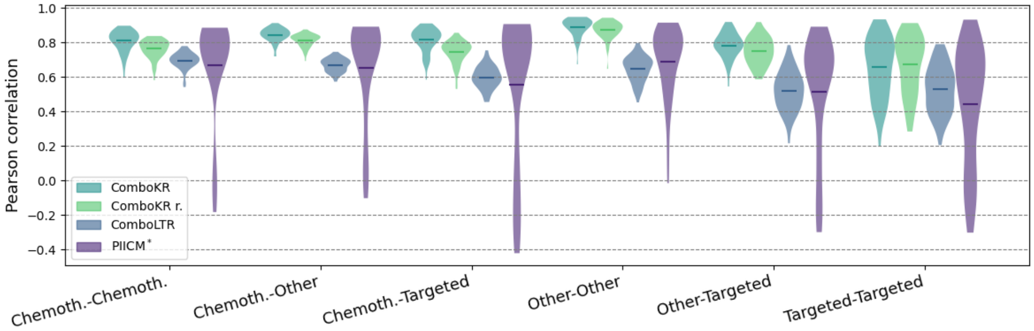


(d) New drug experimental scenario, drugtype combinations

Figure 6: The distributions of Pearson correlations of the drug-dose response prediction on the different tissue and drug type combinations in the two predictive scenarios. The vertical lines in the plots highlight the mean. When fewer than ten samples, the individual results are displayed with crosses.

Table 6: Results of the two-sample Kolmogorov–Smirnov test on the different methods, on the response predictions as well as the synergy scores calculated on them for the two predictive scenarios. p-values less than 0.01 are highlighted with light grey, and those less than 0.05 with dark grey.

(a) Responses, new drug

| | ComboKR | ComboKR r. | ComboLTR |
|---|---|---|---|
| ComboKR r. | 1.16e-07 | | 2.16e-22 |
| ComboLTR | 1.70e-28 | 2.16e-22 | |
| PIICM* | 4.63e-03 | 8.70e-03 | 2.90e-12 |

(b) Responses, new combo

| | ComboKR | ComboKR r. | ComboLTR |
|---|---|---|---|
| ComboKR r. | 9.87e-01 | | 4.76e-05 |
| ComboLTR | 1.12e-04 | 4.76e-05 | |
| PIICM* | 3.78e-01 | 3.78e-01 | 3.51e-07 |

(c) Bliss, new drug

| | ComboKR | ComboKR r. | ComboLTR |
|---|---|---|---|
| ComboKR r. | 3.16e-09 | | 2.67e-01 |
| ComboLTR | 1.16e-07 | 2.67e-01 | |
| PIICM* | 8.70e-03 | 2.37e-03 | 4.63e-03 |

(d) Bliss, new combo

| | ComboKR | ComboKR r. | ComboLTR |
|---|---|---|---|
| ComboKR r. | 9.99e-01 | | 1.10e-08 |
| ComboLTR | 8.64e-10 | 1.10e-08 | |
| PIICM* | 1.20e-01 | 1.20e-01 | 1.12e-04 |

(e) Loewe, new drug

| | ComboKR | ComboKR r. | ComboLTR |
|---|---|---|---|
| ComboKR r. | 3.16e-09 | | 4.67e-02 |
| ComboLTR | 5.57e-11 | 4.67e-02 | |
| PIICM* | 8.70e-03 | 2.37e-03 | 1.17e-03 |

(f) Loewe, new combo

| | ComboKR | ComboKR r. | ComboLTR |
|---|---|---|---|
| ComboKR r. | 9.28e-01 | | 2.25e-14 |
| ComboLTR | 6.10e-13 | 2.25e-14 | |
| PIICM* | 1.82e-01 | 1.20e-01 | 3.16e-09 |
